# Pain-free resting-state functional brain connectivity predicts individual pain sensitivity

**DOI:** 10.1101/790709

**Authors:** Tamas Spisak, Balint Kincses, Frederik Schlitt, Matthias Zunhammer, Tobias Schmidt-Wilcke, Zsigmond T. Kincses, Ulrike Bingel

## Abstract

Individual differences in pain perception are of key interest in basic and clinical research as altered pain sensitivity is both a characteristic and a risk factor for many pain conditions. It is, however, unclear how individual susceptibility to pain is reflected in the pain-free resting-state brain activity and functional connectivity.

Here, we identified and validated a network pattern in the pain-free resting-state functional brain connectome that is predictive of interindividual differences in pain sensitivity. Our predictive network signature (https://spisakt.github.io/RPN-signature) allows assessing the individual susceptibility to pain without applying any painful stimulation, as might be valuable in patients where reliable behavioural pain reports cannot be obtained. Additionally, as a direct, non-invasive readout of the *supraspinal neural contribution* to pain sensitivity, it may have broad implications for translational research and the development and assessment of analgesic treatment strategies.

## Introduction

Pain is a subjective, unpleasant sensory and emotional experience^1^ that is highly variable across individuals^2^. Individual differences in pain perception are of key interest in clinical practice as altered pain sensitivity is both a characteristic and risk factor for many pain conditions^2–4^. In the past decades, brain imaging has revealed the richness and complexity of brain activity underlying both the acute pain experience^5,6^ and pain sensitivity^7,8^. Still, the central nervous mechanisms determining individual differences in pain perception are poorly understood, partly because past neuroimaging research has mainly focussed on the *momentary* (acute or chronic) pain experience. The common practice of using pain-free episodes merely as a baseline reference makes studies inherently blind to components of brain activity that are not time-locked to painful events but still central to pain processing and perception.

Brain activity in the ‘resting-state’ (i.e. in absence of any task or stimulation) is known to mirror some, if not all, task-induced activity patterns^9^. For instance, the well-known ‘large-scale resting-state networks’^10^ (RSNs) strongly resemble patterns of related tasks^11^. Moreover, resting state activity can predict behavioural performance, perceptual decisions and related neural activity^12,13^. Given the tight link between resting-state and task-induced brain activity, it is highly plausible that activity and functional connectivity during *pain-free* resting-state conditions reflect the individual’s susceptibility to pain. Following the RSN-related terminology, we refer to this type of neural activity as the ‘*resting-state network of pain susceptibility*’.

This proposed pain-related resting-state network might have been captured by studies reporting that brain activity and connectivity directly preceding pain^14–21^ and the variability of resting-state fMRI signal^22^ in pain-free state is associated with the neural response to nociception and the resultant pain experience. However, due to the small sample sizes, highly varying methodology (e.g. regarding the correction of physiological and motion artefacts) and the lack of validation in these previous studies (‘replication crisis’^23^), the predictive power and clinical relevance of this kind of resting-state brain activity remains unclear to date^24^.

Mapping the *resting-state network of pain susceptibility* and exploiting its capacity to predict various aspects of pain processing would substantially advance the field – both from a basic research and translational perspective. First, contrasting it with experimental pain responses would extend our understanding of how the subjective experience of pain emerges from brain activity. Second, investigating how the hypothesised *resting-state pain susceptibility network* is embedded into the broader resting-state brain activity could extend our knowledge about the complex functional architecture of the resting brain.

Finally, and most importantly, a robust, generalizable and rigorously validated prediction of pain sensitivity - based on the ‘*resting-state network of pain susceptibility’ –* could lay the foundations for a non-invasive neuromarker of an individual’s sensitivity to pain. Such a resting brain network-based biomarker could be an essential part of a future ‘pain biomarker composite signature’^24^ required for diagnosis and for evaluation of risk of developing pain and of analgesic efficacy and objectively characterise pain conditions and analgesic treatment effects in experimental and clinical pain research.

Here, we investigated the capacity of *pain-free* resting-state functional brain connectivity to predict individual pain sensitivity in a sample of a total of *N=116* young healthy participants. We first performed a whole brain search for specific *features* of the *pain-free* resting-state connectome, which are predictive for individual pain sensitivity in a sub-study used only for the ‘*training’* and ‘*internal validation’ of the predictive model*. Then, we performed a *prospective* validation of the approach in terms of predictive performance, generalisation and potential confounders in two independent sub-studies acquired at different scanning sites (‘*external validation’*). Finally, we performed a *‘reverse-mapping’* of the predictive model to identify the *key nodes* of the hypothesised network, hereinafter referred to as the ‘*Signature of the Resting-state Pain-susceptibility Network*’ (abbreviated as RPN-signature).

## Results

### Functional connectivity-based prediction and independent multi-centre validation

Resting-state functional MRI data were obtained from a total of *N=116* healthy volunteers over three separate sub-studies, performed in three different imaging centres. Neuromarker development was based on intrinsic whole-brain functional brain connectivity, the degree to which resting-state brain activity in distinct neural regions is correlated over time (in the absence of any explicit task). Direct functional connectivity was assessed between *M=122* functionally defined regions (Figure **1**). Heat, cold and mechanical pain thresholds obtained according to the well-established quantitative sensory testing (QST) protocol^25^ were aggregated in a composite pain-sensitivity score, as previously reported^8^. Whole-brain resting-state functional connectivity data of *study 1* (*N*_*1*_*=35*, after exclusions) was used as the input feature-space (*P=7503* features per participant) to predict individual pain sensitivity scores, leading to a typical “large *P* — small *N*” setting.

**Figure 1.**
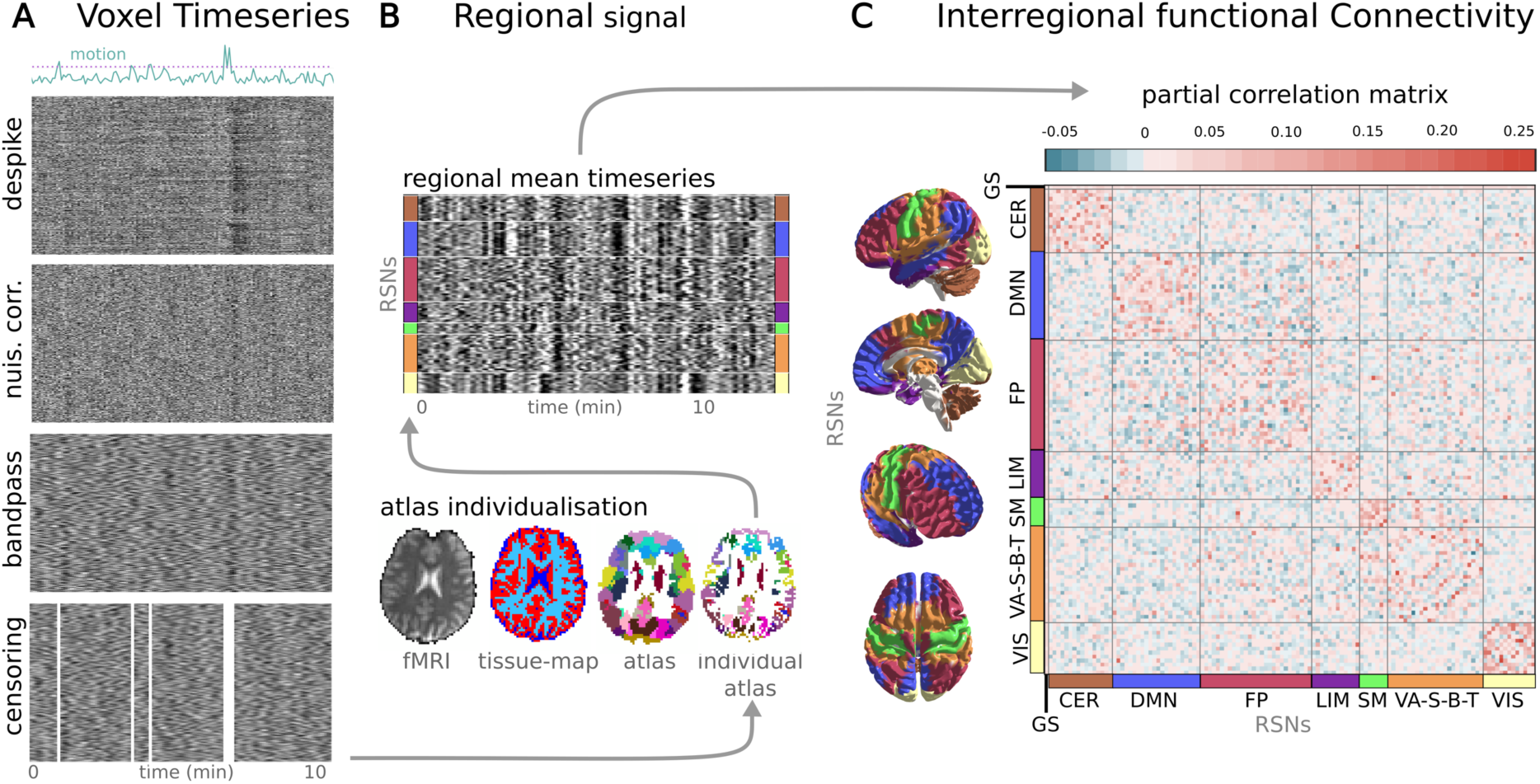
Calculating functional brain connectivity from resting state fMRI measurements. Raw brain images of *N=116* participants, in total, underwent automated artefact removal, involving despiking, nuisance regression, bandpass filtering and censoring of motion contaminated time-frames. The effects of these procedures on the BOLD signal are exemplified on the carpet-plots (**A**, x: time, y: voxel, colour: intensity). Subsequently, a multi-stage, high-precision brain atlas individualisation was performed to obtain regional grey-matter signals for *M=122* functionally defined brain regions (**B**). Partial correlation between all possible region pairs was computed to asses direct functional connectivity and ordered based on large-scale modularity to form individual connectivity matrices. Partial correlations of all regions with the global grey matter signal was retained to account for, but not completely discard the effect of the “global signal”, a component of brain activity often regarded as a confound but also related to e.g. vigilance^26^. (**C**) Subject-level connectivity matrices (depicted by the group-mean connectivity matrix) from Study 1 served as an input for machine-learning-based prediction of behavioural pain sensitivity. The functional connectivity workflow, together with the predictive model, is open-source and shared with the community as a docker container (*https://spisakt.github.io/RPN-signature*).

According to these conditions, we constructed a machine-learning pipeline, consisting of feature-normalisation, feature-selection and fitting an elastic net regression model. Model training consisted of fitting the pipeline and optimising its hyperparameters in a leave-one-participant-out cross-validation framework to improve generalisation to new data.

In *Study 1*, QST-based pain sensitivity values ranged from *-1.45* to *1.52* with a robust range (range between the 5^th^ and 95^th^ percentiles) of *2.57* (in arbitrary units).

Internal validation (i.e. performance on left-out participant data) revealed that the model predicts pain sensitivity with a mean absolute error of *MAE*_*1*_*=0.56* (mean squared error: MSE_1_=0.32, median absolute error: MedAE_1_=0.27, *Explained variance Expl.Var.*_*1*_*=*39%, *Pearson’s r*_*1*_*=0.63, p*_*1*_*=0.00006*, Figure 2B). Diagnostics of the model fit (learning curve analysis, Figure 2A) suggested that the approach reduced overfitting and that sample size was sufficient for an acceptable generalisation. The machine-learning pipeline with the optimal hyperparameters was finally fitted to the data of all participants in *Study 1* and saved for further use. The model trained in *Study 1* is henceforth referred to as the signature of the Resting-state Pain-sensitivity Network (or short, **RPN signature**).

**Figure 2.**
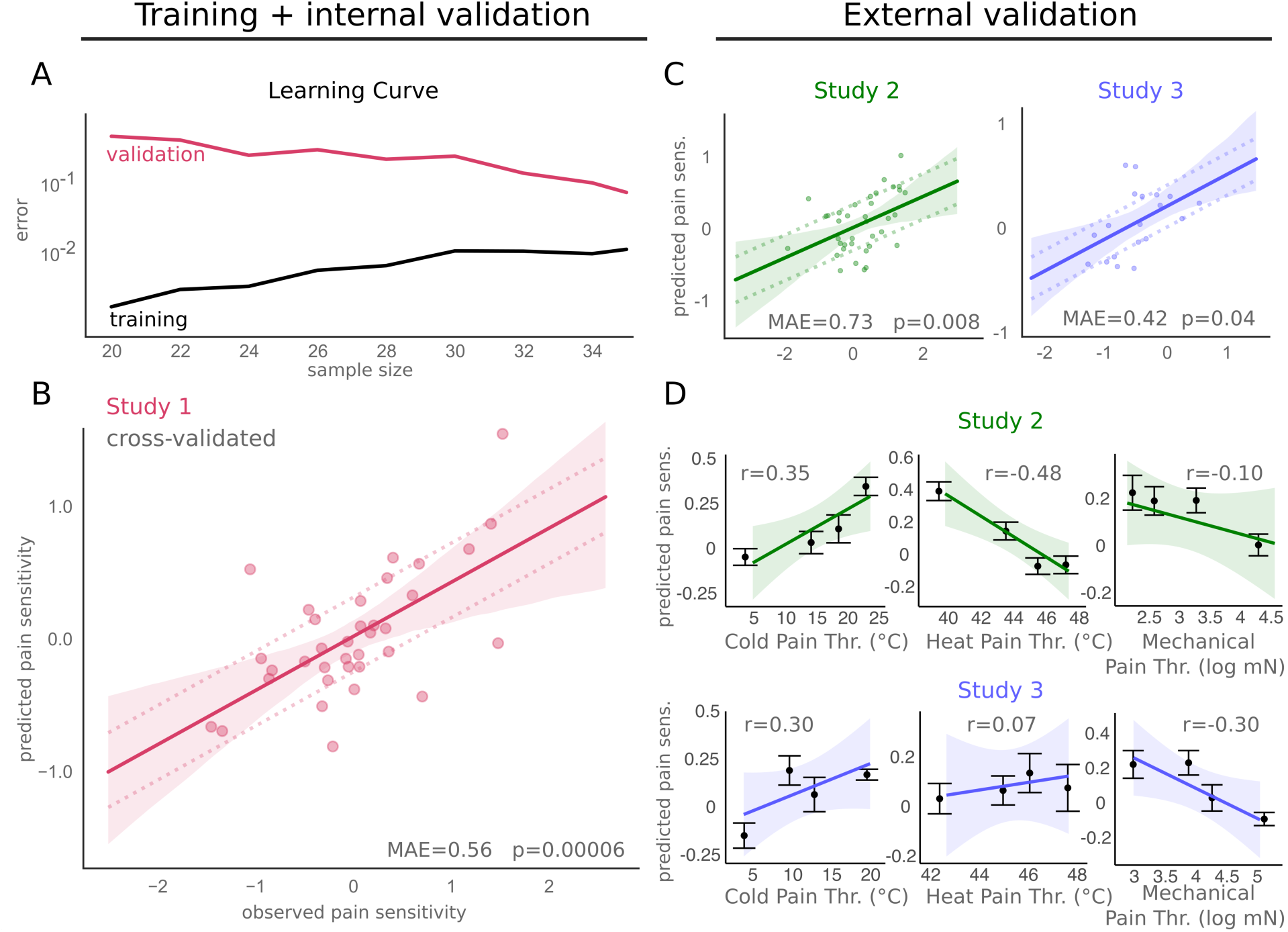
The RPN-signature predicts individual pain sensitivity based on pain-free resting-state functional brain connectivity. The learning-curve (**A**) suggests that the size of the training sample (*Study 1*) was sufficient to substantially reduce overfitting and improve generalisation. Internal cross-validated prediction in the training sample (**B**) and prospective external validation in the test samples (*Studies 2* and *3*, **C**) revealed considerable predictive accuracy, robustness, and multi-centre generalisability of the RPN-signature. Mean absolute error (MAE) of the prediction is depicted by dashed lines. While the prediction target was the composite pain sensitivity score, the predicted values correlate with its sub-components, the QST sensory modalities (cold, heat and mechanical pain thresholds, **D**), as well. Shaded ribbons imply the 95% confidence intervals for the regression estimates. On panel **D**, points represent the means values of quartile-based sub-groups and bars represent standard errors.

The full pipeline is deployed as a BIDS-app, a software tool following the Brain Imaging Data Structure (BIDS) specification^27^ (https://spisakt.github.io/RPN-signature).

As pre-registered on the 7^th^ of March, 2018 (http://osf.io/buqt7), external validation studies (*Studies 2* and *3*) were performed in different imaging centres with different MRI scanners (from three different vendors) and with different research staff.

The multi-centre design, together with a reasonable variability in imaging sequences (see *Table 2* of the on-line methods) introduced an inherent heterogeneity ensuring that test samples are maximally independent and provide a realistic estimate of prediction accuracy and generalisability.

In *Studies 2* and *3* (*N*_*2*_*=37* and *N*_*3*_*=19*, after exclusions), QST-based pain sensitivity values ranged from *-1.82* to *1.57* and from *-1.2* to *0.55*, with a robust range of *2.3* and *1.43*, respectively (in arbitrary units). External validation (Figure 2C) revealed a considerable generalisability of the predictive model: the mean absolute prediction error was *MEA*_*2*_*=0.73* and *MAE*_*3*_*=0.42*, respectively (*MedAE*_*2*_*=0.55, MedAE*_*3*_*=0.25, MSE*_*2*_*=0.54, MSE*_*3*_*=0.17, Expl.Var.*_*2*_*=18%, Expl.Var.*_*3*_*=17%, Pearson’s r*_*2*_*=0.43* and *r*_*3*_*=0.47, p*_*2*_*=0.008* and *p*_*3*_*=0.04*). The predicted pain sensitivity score was not only correlated with the observed QST-based composite pain sensitivity score but also with its modality-specific components, the cold, heat and mechanical pain thresholds (*CPT, HPT* and *MPT*, respectively) in both *Study 2* (*r*_*CPT,2*_*=0.35, r*_*HPT,2*_*=0.48, r*_*MPT,2*_*=0.10)* and *Study 3* (*r*_*CPT,3*_*=0.30, r*_*HPT,3*_*=0.07, r*_*MPT,3*_*=0.30*) (Figure 2D). Summary statistics of pain sensitivity, in-scanner motion and demographic data are reported in Table S1.

### Potential confounds and specificity to pain sensitivity

To ensure that the RPN-signature captures the pain-related neural processing in the pain-free resting state, the potential contribution of two types of confounds has to be ruled out: *(i)* imaging artefacts (e.g. head motion artefacts^26^) and *(ii)* demographic or behavioural variables correlated with individual pain sensitivity (e.g. age and sex are known to be slightly correlated with QST thresholds^25^).

Table 1 lists the investigated (pre-registered) confounder variables and their correlations to the predicted pain sensitivity score (together with the corresponding p-value) in all three studies. The pain sensitivity score predicted by the RPN-signature was *not* significantly associated with any of the confounder variables (p>0.05 for all variables). Effect size was, however, considerable for sex (*Study 2*: *R*^*2*^*=0.08, p=0.09*), number of days from the first day of menses (*Study 1*: *R*^*2*^*=0.26, p=0.11, Study 2*: *R*^*2*^*=0.11, p=0.17, Study 3*: *R*^*2*^*=0.33, p=0.08*), time difference between the MRI and QST measurements (*Study 2: R*^*2*^*=0.1, p=0.06*) and with Glutamate/Glutamine levels in pain processing regions (measured by MR spectroscopy in *Study 1*: *R*^*2*^*=0.09, p=0.08*). Summary statistics of confounder variables are reported in Table S1.

**Table 1.** Confounder analysis: The RPN signature-response is specific to pain sensitivity. No significant association (p < 0.05) was found with any of the confounder variables. Effect sizes with R^2^ ≥ 0.09 (medium effect size according to Cohen^28^) and p-values less than 0.1 is denoted by bold letters. In Study 1, MRI and QST was performed on the same day, otherwise it was measured within 1-5 days. GABA and glutamine/glutamate levels were measured by MR spectroscopy in regions of the pain matrix^8^. Abbreviations: FD: framewise displacement, % scrubbed: number of censored time-frames, BP: blood pressure, mens. day: number of days since the first day of menstruation on the MRI-day, catas.: catastrophising, sensitiv.: sensitivity, CDT, WDT, MDT: cold, warmth and mechanical detection thresholds, T50: temperature inducing moderate pain. BP is reported here as measured on the day of the MRI measurement. See Table S3 for additional covariates (trait anxiety and BP on the day of QST measurement, all p>0.1).

| MRI<br>confounds | median FD |  | max FD |  | % scrubbed |  | BP sys. |  | BP dias. |  | MRI-QST<br>dif. |  |
| --- | --- | --- | --- | --- | --- | --- | --- | --- | --- | --- | --- | --- |
|  | R<sup>2</sup> | p | R<sup>2</sup> | p | R<sup>2</sup> | p | R<sup>2</sup> | p | R<sup>2</sup> | p |  |  |
| **Study 1** | 0.014 | 0.506 | 0.004 | 0.705 | 0.008 | 0.613 | N/A | N/A | N/A | N/A | - | - |
| **Study 2** | 0.026 | 0.339 | 0.030 | 0.304 | 0.021 | 0.388 | 0.034 | 0.289 | 0.026 | 0.352 | **0.099** | **0.058** |
| **Study 3** | 0.006 | 0.745 | 0.025 | 0.517 | 0.057 | 0.324 | 0.000 | 0.943 | 0.000 | 0.951 | 0.001 | 0.880 |
| demography | age |  | sex |  | BMI |  | mens. day |  | alcohol |  | education |  |
|  | R<sup>2</sup> | p | R<sup>2</sup> | p | R<sup>2</sup> | p | R<sup>2</sup> | p | R<sup>2</sup> | p | R<sup>2</sup> | p |
| **Study 1** | 0.010 | 0.572 | 0.000 | 0.943 | 0.012 | 0.511 | **0.26** | 0.107 | 0.027 | 0.343 | 0.012 | 0.537 |
| **Study 2** | 0.001 | 0.840 | 0.081 | **0.089** | 0.010 | 0.555 | **0.11** | 0.173 | 0.000 | 0.913 | 0.038 | 0.252 |
| **Study 3** | 0.080 | 0.240 | 0.010 | 0.677 | 0.000 | 0.955 | **0.33** | **0.078** | N/A | N/A | N/A | N/A |
| psychometrics | pain catas. |  | pain sensitiv. |  | depression |  | sleep quality |  | anxiety state |  | stress |  |
|  | R<sup>2</sup> | p | R<sup>2</sup> | p | R<sup>2</sup> | p | R<sup>2</sup> | p | R<sup>2</sup> | p | R<sup>2</sup> | p |
| **Study 1** | 0.000 | 0.958 | 0.011 | 0.551 | 0.002 | 0.789 | N/A | N/A | 0.002 | 0.789 | N/A | N/A |
| **Study 2** | 0.062 | 0.136 | 0.017 | 0.437 | 0.021 | 0.390 | 0.008 | 0.600 | 0.021 | 0.390 | 0.003 | 0.730 |
| **Study 3** | 0.012 | 0.657 | 0.005 | 0.773 | **0.094** | 0.216 | 0.029 | 0.547 | **0.094** | 0.216 | **0.127** | 0.134 |
| QST and<br>MRS | CDT |  | WDT |  | MDT |  | T50 |  | GABA |  | GLX |  |
|  | R<sup>2</sup> | p | R<sup>2</sup> | p | R<sup>2</sup> | p | R<sup>2</sup> | p | R<sup>2</sup> | p | R<sup>2</sup> | p |
| **Study 1** | 0.002 | 0.791 | 0.002 | 0.786 | N/A | N/A | 0.039 | 0.255 | 0.048 | 0.205 | **0.089** | **0.081** |
| **Study 2** | 0.014 | 0.487 | 0.000 | 0.957 | 0.041 | 0.230 | N/A | N/A | N/A | N/A | N/A | N/A |
| **Study 3** | 0.019 | 0.579 | 0.055 | 0.334 | 0.027 | 0.502 | N/A | N/A | N/A | N/A | N/A | N/A |

### The RPN-signature: the predictive resting-state network of the pain sensitivity

In the applied machine-learning pipeline, non-zero regression coefficients naturally delineate the predictive sub-network. Each coefficient can be interpreted as the relative “importance” of the connectivity in the prediction. Positive (negative) coefficients translate to “stronger interregional functional connectivity predicted higher (lower) sensitivity to pain”.

The RPN-signature model, trained in *Study 1*, retained 21 non-zero links out of the total number of 7503 functional connections. The predictive connections are listed in Table 2 and the predictive network is depicted on the chord plot of Figure 3B.

**Table 2.**
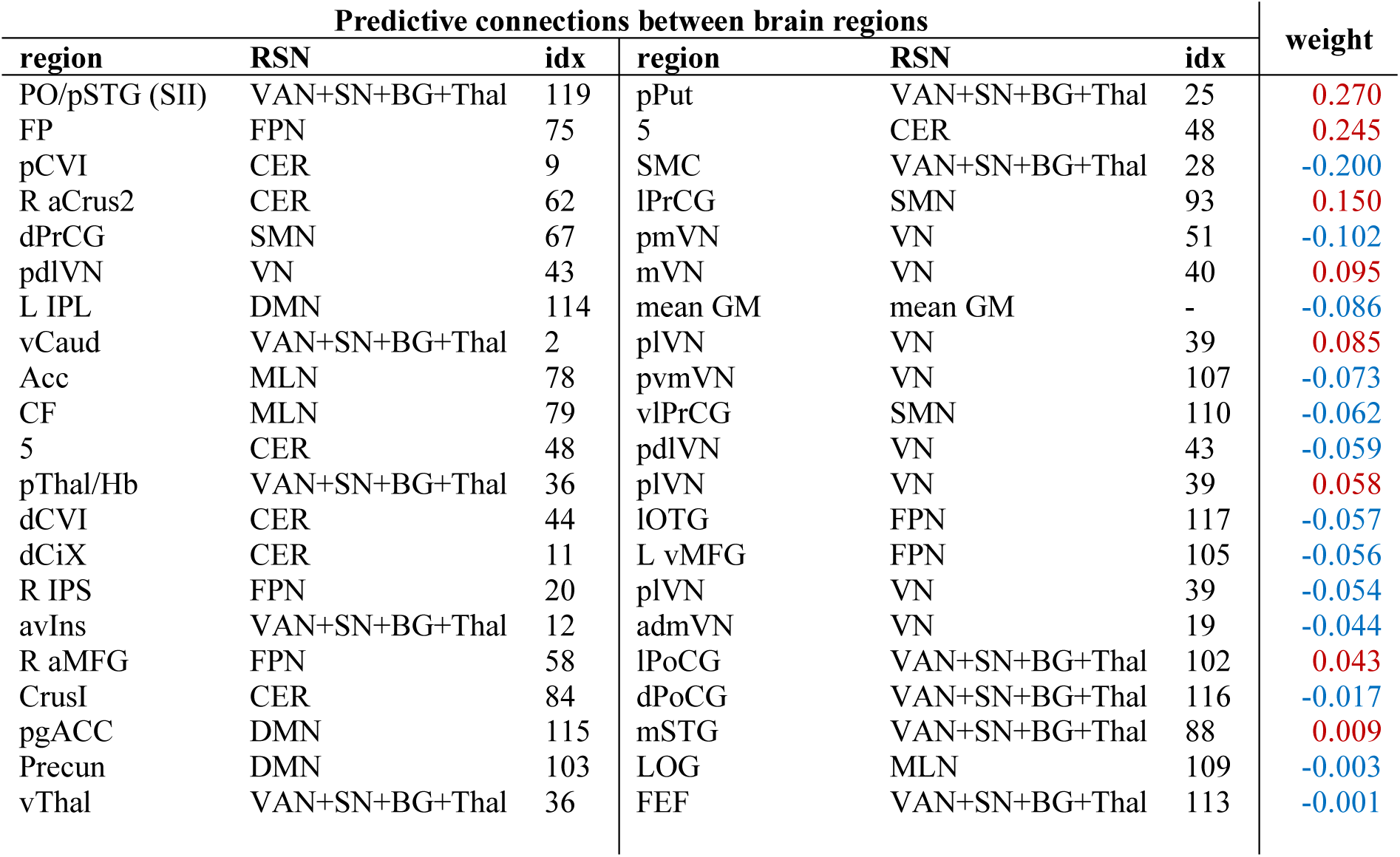
Predictive connections of the RPN signature. Non-zero regression coefficients naturally delineate the predictive sub-network. Regions and corresponding large-scale resting-state network (RSN) modules are to be interpreted as in the MIST atlas (see Methods, original atlas-index is given). Predictive connections are ordered by their absolute predictive weights. Connectivity strengths associated with higher and lower sensitivity to pain are highlighted in red and blue, respectively. Abbreviations: CER: cerebellum, roman numbers: cerebellar lobes, GM grey matter, VAN: ventral attention network, SN: salience network, BG: basal ganglia, Thal: thalamus, Hb: habenula, MLN: mesolimbic network, FPN: frontoparietal network, SMN: somatomotor network, DMN: default mode network, VN: visual network, Ins: insula, PO: parietal operculum, SII: secondary somatosensory cortex, STG: superior temporal gyrus, FEF: frontal eye-field, PrCG: precentral gyrus, PoCG: postcentral gyrus, SMC: supplementary motor cortex, Put: putamen, Caud: caudate nucleus, Acc: nucleus accumbens, LOG: lateral orbital gyrus, CF: collateral fissure, OTG: occipitotemporal gyrus, MFG: middle frontal gyrus, IPS: intraparietal sulcus, pgACC: perigenual anterior cingulate cortex, PrC: precuneus cortex. Prefix: L: left, R: right, a: anterior, p: posterior, v: ventral, d: dorsal, l: lateral, m: medial.

**Figure 3.**
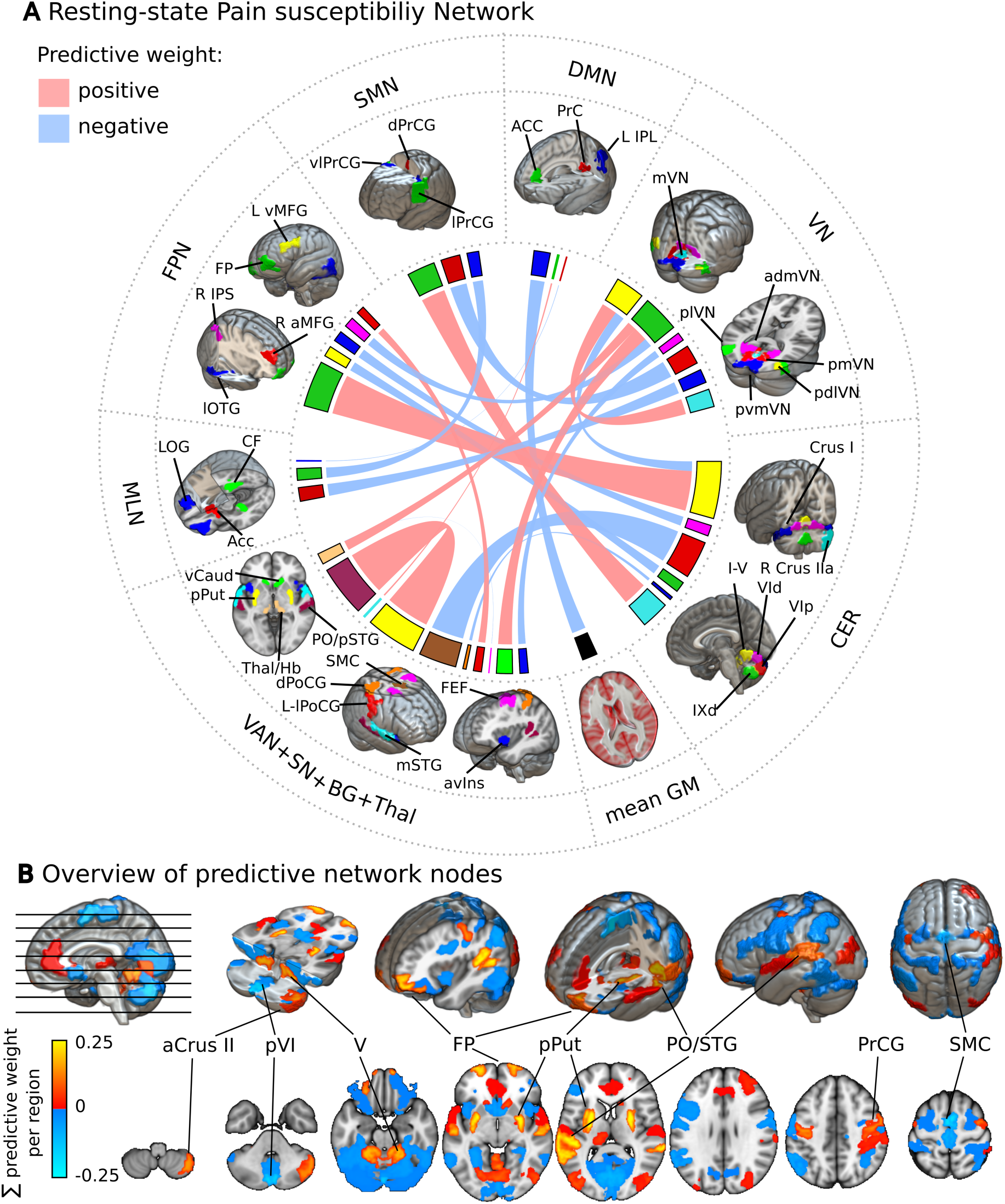
The resting-state pain sensitivity network signature. **(A)** The predictive network of the RPN-signature. Widths of ribbons are proportional to the predictive weights of the functional connections. Network-nodes are color-coded and displayed in 3D-views. Note that, the utilised brain atlas is based on an entirely data-driven functional parcellation and is, therefore, not fully bilateral. Where laterality (L: left, R: right) is not explicitly specified, the atlas did not distinguish the region from its contralateral homolog. **(B)** Regional predictive strength map of the RPN-signature. Colour-bar depicts ‘region-wise predictive strength’ (sum of the weights of all connections for the region, multiplied by the study-specific regional probability map). Regions with an absolute predictive strength greater than 0.1 are annotated. Abbreviations: CER: cerebellum, roman numbers: cerebellar lobes, GM grey matter, VAN: ventral attention network, SN: salience network, BG: basal ganglia, Thal: thalamus, Hb: habenula, MLN: mesolimbic network, FPN frontoparietal network, SMN: somatomotor network, DMN: default mode network, VN: visual network, Ins: insula, PO: parietal operculum, SII: secondary somatosensory cortex, STG: superior temporal gyrus, FEF: frontal eye-field, PrCG: precentral gyrus, PoCG: postcentral gyrus, SMC: supplementary motor cortex, Put: putamen, Caud: caudate nucleus, Acc: nucleus accumbens, LOG: lateral orbital gyrus, CF: collateral fissure, OTG: occipitotemporal gyrus, MFG: middle frontal gyrus, IPS: intraparietal sulcus, pgACC: perigenual anterior cingulate cortex, PrC: precuneus cortex. Prefix: L: left, R: right, a: anterior, p: posterior, v: ventral, d: dorsal, l: lateral, m: medial.

Almost half of the variance in the predicted pain sensitivity score is explained by the 4 “strongest” connections. The most important *positive* predictive connections are found between: *(i)* the posterior putamen (pPut) and a region including parts of the parietal operculum and the posterior superior temporal gyrus (PO/pSTG); *(ii)* the frontal poles (FP) and the cerebellar lobule V; and *(iii)* the right anterior crus II of the cerebellum and the lateral precentral gyrus (lPrCG, primary motor cortex). The only *negative* predictor among the ‘top 4’ connections was a connection between the supplementary motor cortex and the posterior part of cerebellar lobule VI. Several other interregional connections and, additionally, the global grey matter signal was also found contribute to the predicted pain sensitivity.

To simplify the overview of the spatial pattern of regions involved in the RPN-signature, we calculated the node-wise sums of predictive weights and multiplied it with the study-specific regional probability-maps. The resulting node-wise predictive strength map is displayed on Figure 3A. To aid the interpretation of our results, we contrasted the localisation of the network-nodes in the RPN-signature (Figure 4C) to two voxel-level multivariate pain-signatures, the Neurologic Pain Signature (NPS, Figure 4A) and the “extra-nociceptive”, stimulus independent contribution to pain processing, as represented by the Stimulus Intensity Independent Pain Signature-1^29^ (SIIPS1, Figure 4B).

**Figure 4.**
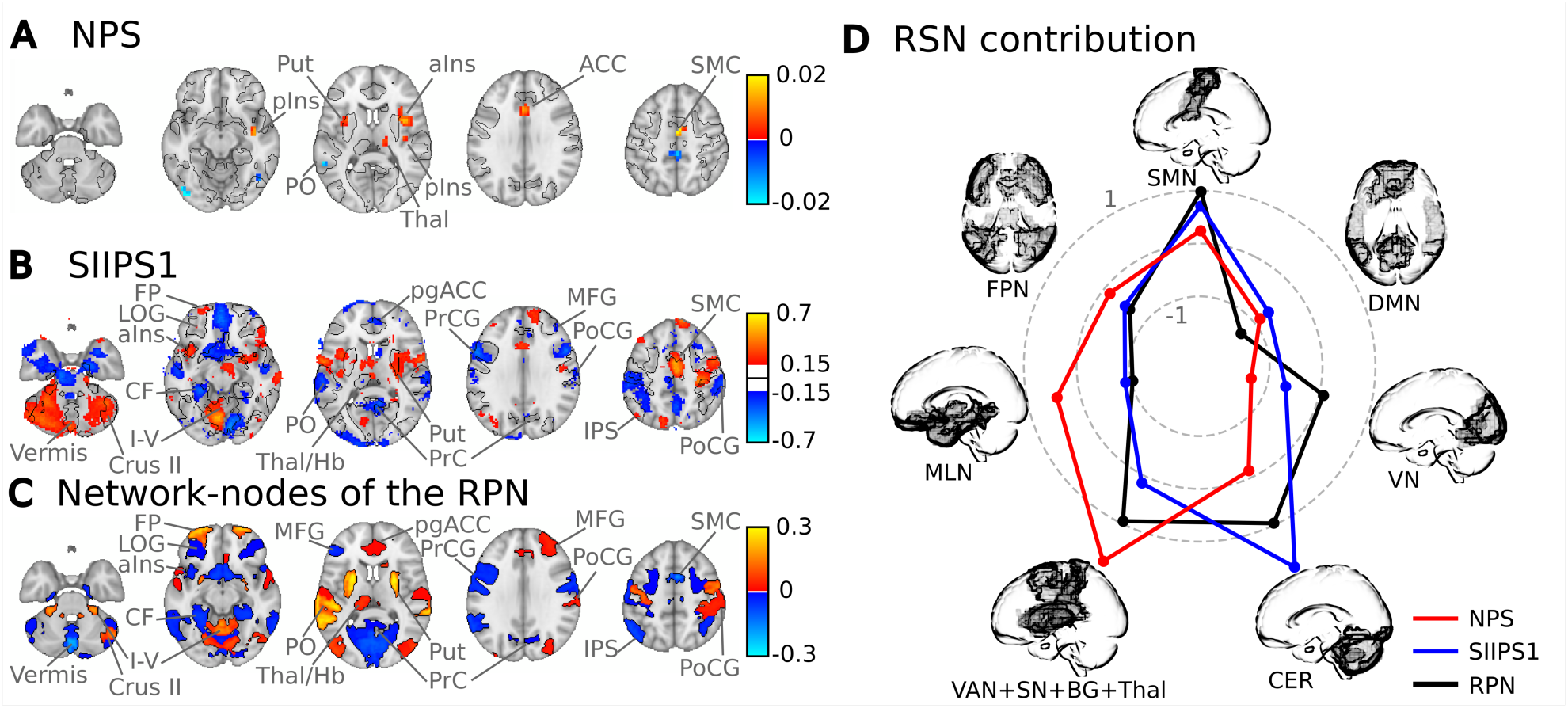
Overlap of the network-nodes in the RPN signature with voxel-level multivariate predictive signatures of pain. The anterior insula (aIns), thalamus (Thal), parietal operculum (PO), putamen (Put), anterior cingulate (ACC) and supplementary motor (SMC) cortices, where pain-elicited activation is predictive of the intensity of pain, as mapped by the Neurologic Pain Signature (NPS, **A**)^30^, are also network-nodes in the RPN-signature (**C** and black outlines on **A, B** and **C**). The overlap with pain-regions is even more striking if the effect of stimulus intensity is ruled out, as in the case of stimulus intensity independent pain signature-1 (SIIPS1, **B**)^29^. The pain-elicited activity of the lateral orbital gyrus (LOG), left middle frontal gyrus, (MFG), perigenual anterior cingulate cortex (pgACC), precentral gyrus (PrCG), parietal operculum (PO), postcentral gyrus (PoCG) and several cerebellar regions (mostly the crus II and vermis VIII) is predictive to stimulus-intensity independent pain experience and their resting-state functional connectivity predicts individual pain sensitivity. Focusing on the involvement of the large-scale functional brain networks in the investigated predictive patterns **(D)** reveals that the activity/connectivity of the ventral attention network (VAN), salience network (SN), basal ganglia (BG), thalamus (Thal) and somatomotor network is prominent in all investigated predictive signatures. Moreover, while the NPS involves the mesolimbic network, cerebellar activity/connectivity strongly affects the SIIPS1 and the RPN signatures. Note that, while the spatial patterns of the signatures show a considerable overlap, the voxel-level activity-based signatures (NPS and SIIPS1) are conceptually different from the network-level RPN-signature, therefore a direct comparison of the magnitude and sign of the predictive weights and regional predictive connectivity strengths is not possible.

There is a striking overlap between the RPN network nodes and voxel-level multivariate pain-signatures, involving the aIns, pgACC, PO, pPut, SMC, thalamus (Thal, posterior part, also including the habenula, Hb) and in case of the SIIPS1, also the FP, PrCG, PoCG, PrC, MFG (dlPFC), LOG (vlPFC) and the cerebellar crus II, lobules I-V and vermal regions. Note that the signs of the predictive weights of the NPS and the SIIPS1 represent brain activity and it is, therefore, not directly comparable to the RPN node-wise predictive connectivity weights.

Analysing the contribution of the seven large-scale resting-state networks (RSNs from the MIST brain atlas, Figure 4D) to these patterns revealed that the functional conglomerate consisting of the ventral attention and salience networks, the basal ganglia and the thalamus (VAN+SN+BG+Thal) and, additionally, the CER and the SMN were highly involved in all three patterns. The NPS exhibited the strongest relative VAN+SN+BG+Thal network activation. Moreover, it displayed a stronger MLN and FPN and a lower visual network (VN) and CER contribution than the SIIPS1. The RSN-wise pattern of the RPN-signature was similar to both of the NPS and the SIIPS1, but seems to be closer to the Stimulus Intensity Independent Signature of pain processing.

## Discussion

Here we report the RPN-signature, an objective, *brain-based* measure of pain sensitivity, based on *functional connectivity* acquired during *pain-free* resting-state. The applied prospective validation procedure establishes solid foundations for promising basic research and translational applications. The RPN-signature is to be applied together with a fixed resting-state fMRI analysis pipeline (freely available as source-code and *Docker-based BIDS-application* at *https://spisakt.github.io/RPN-signature*) and provides the opportunity for *out-of-the-box* resting-state fMRI-based, non-invasive characterisation of pain sensitivity.

This work addresses an important gap in basic research by providing strong evidence for the association of *pain-free* resting-state functional brain connectivity with neural processing of painful stimuli and the corollary pain experience. The identified functional network pattern provides novel insights into this - commonly unaccounted - component of resting-state brain activity and substantially advances our understanding of the neural mechanisms underlying an individual’s pain sensitivity. We used a pre-registered, multicentre design and deployed a substantial sample size to perform a rigorous *prospective validation* of our predictive model. Therefore, the RPN-signature may serve as an objective neuromarker of interindividual differences and alterations in pain sensitivity^2–4^.

It is important to distinguish our study of the “pain-free” resting-state from other predictive efforts in pain research, like patient-control classification studies^31^ (but also “pain decoding”^30^), which examine brain activity in experimental or chronic pain conditions, i.e. in the presence of painful experience. Note, that in studies of chronic pain, the terminology “resting-state” usually refers to the lack of explicit experimental pain stimuli in the data acquisition paradigm, but not to the lack of ongoing spontaneous pain experience.

In contrast, the RPN-signature is based on brain activity measured in the absence of any ongoing painful experience (which we refer to as “pain-free” resting-state). Therefore, it introduces a conceptually new modality for future efforts of building a composite pain biomarker^24^.

### Predictive power and clinical relevance of the RPN-signature

The RPN-signature predicts a considerable amount of the variance in individual pain sensitivity (*39%* with internal validation and *18%-19%* with external validation, Figure 2) which, according to Cohen’s recommendations^28^, can be considered as being in the ‘*medium-to-large*’ range. The mean absolute error (MAE) of the prediction was *0.73* and *0.42* in the external validation studies (and *0.56* with internal validation). Interpreting the magnitude of the error in comparison to the min-max-range (3.39) and the inter-quartile-range (1.04) of the observed pain sensitivity values strongly suggests that the predictive power of the RPN-signature is clinically relevant in the context of chronic pain^32^ and renders the RPN signature deployable in numerous applications. Here, we discuss three aspects of evaluating the relevance of the achieved predictive performance.

First, the prediction accuracy we report is similar to those in previous resting state fMRI studies of other target variables (see e.g. Table A2 of Ref.^33^). However, we would like to point out the majority of previously used fMRI-based predictive models were only internally validated (i.e. the same dataset was used for model training and validation) whereas our validation is based on two independently acquired samples at different scanning sites. The level of prediction accuracy we reached in this more rigorous, prospective approach therefore underscores the robustness of the findings.

Second, our study is based on a large sample size (*N=116*) and the external validation featured heterogeneity regarding methodology, infrastructure, research personnel and a 1-5-day delay between the QST and the MRI measurements (which were at the same day in the training dataset, see Table S1 for further details). Our study thus overcomes recent concerns about common methodological pitfalls of neuroimaging based predictors^34,35^. While prediction accuracy estimates could likely be higher with a stricter standardisation of the research protocols, the above reported estimates are expected to robustly generalise to a wider variety of resting-state fMRI datasets.

Third, we believe that relying solely on the QST-based predictive accuracy might lead to underestimating the utility of the RPN-signature. While Quantitative Sensory Testing is the gold standard approach to assess pain sensitivity, it is a measure of subjective experience, shaped by *peripheral, spinal* and *supraspinal* processes convolved with perceptual and behavioural error components^36^. The RPN-signature, on the other hand captures signal of *supraspinal* neural origin only. Due to this difference, prediction accuracy estimates based on the ‘multifaceted’ QST-based observations should serve only as a *lower bound* when judging the utility of the RPN-signature as a proxy for measuring the *supraspinal neural component* of the interindividual variability of pain sensitivity.

### Neuroscientific validity and specificity of the RPN-signature

The predicted pain sensitivity is not only correlated with the observed composite pain sensitivity score but also with its sub-components, the cold, heat and mechanical pain thresholds. The facts that the predicted score shares variance with the sensory modality-specific pain thresholds is remarkable as the predictive model was only informed about the composite score during training. On the other hand, neither the investigated imaging artefacts nor the observed demographic or behavioural variables were significantly correlated with the predicted pain sensitivity values. In sum, our analysis strongly suggests that the predictive power of the RPN-signature is *(i)* based on signal of neural origin, *(ii) is* specific to pain sensitivity and *(iii) is* not driven by the general sensitivity to sensory stimuli or pain-related psychological variables such as anxiety, depression or sleep quality.

QST pain thresholds are known to differ between sexes and in different phases of the menstrual cycle. Their moderate (but statistically not significant) correlations with the predicted pain sensitivity score suggest that the RPN-signature partially captures the neural correlates of these effects. The weak (R^2^=0.09, p=0.08) correlation with Glutamate/Glutamine levels in pain processing regions suggest that the RPN-signature also captures the previously reported^8^ neurotransmitter-level-dependence of individual pain sensitivity. The pain sensitivity scores, predicted by the RPN-signature seem to be also slightly associated to the delay (days) between the MRI and QST measurements, which suggests that pain sensitivity and its resting-state neural correlates are subject to dynamic changes within the scale of days.

### The “Resting-state Pain-sensitivity Network”

The identification of the resting-state connectome that predicts an individual’s pain sensitivity yields important conceptual advances in our understanding of the neural mechanisms underlying pain sensitivity and – likely the susceptibility to pain chronification.

Many of the key hubs of the RPN-signature, such as the PO/SII, pPut, SI, dlPFC, habenula (Hb), pgACC and aIns (Figure 3 and Figure 4) have commonly been associated with pain. The connectivity between the PO/STG area (comprising SII) and pPut displayed the greatest predictive power with a positive association with pain sensitivity. Notably, the PO/STG belongs to the same functional complex as the dorsal posterior insula^37^. The predictive power of this connection corroborates findings suggesting that subregions of the PO^38^ and the dpIns^39^ are specific to nociception and pain related percepts. The involvement of the pPut in pain-related and affective sensorimotor processes is also well known; particularly it shows pain intensity related and partially somatotopic activation during painful stimulation^40,41^. Moreover, a recent finding links grey matter density in pPut and pIns to pain sensitivity^41^. According to current concepts, the multifaceted experience of pain results from the integration of nociception and the cognitive-emotional state of the individual^42^. Within such framework, a stronger resting-state co-activation of the pPut and SII (and dpIns) might imply that the sensory aspects of salient (and possibly nociceptive) inputs have an elevated ‘weight’ in this integrative process.

The integration of cognitive-emotional states into pain perception might be reflected by the activity and connectivity of the dlPFC. Indeed, we found that the connectivity between the postcentral gyrus (SI) and the dlPFC is positively associated with pain sensitivity. This is very much in line with a recent finding^43^, showing that neuromodulation of the dlPFC introduces decreased pain-related activity in sensory-motor areas (and increased activity in the pIns and thalamus) and that this change in activation, together with the resulting analgesic effect, is linked to structural connectivity between the dlPFC and the thalamus.

Modulatory function of both the dlPFC and the caudate-putamen might be projected to the brainstem and particularly to the antinociceptive periaqueductal grey (PAG, not investigated in the present study) by the habenula^44^, which we also found to contribute to the predicted pain sensitivity score. The relevance of this pathway in shaping one’s sensitivity to pain is supported by the previously observed negative correlation between resting-state mPFC-PAG connectivity and intrinsic attention to pain^45^.

In addition to connectivity within the ‘sensory-frontal-striatal-habenula’ complex which explains about 25% of the predicted pain sensitivity score, a second ‘cluster’ of connections also contributes to pain sensitivity. At the centre of this cluster is the cerebellum and its connections to frontal and sensory-motor regions (frontal pole, dlPFC, PrCG, PoCG, SMC). Our finding that the cerebellum significantly shapes individual pain sensitivity corroborates previous reports (see e.g. Ref.^46^ – study on patients with cerebellar infarcts and pain sensitivity and pain modulation by placebo) and highlights the cerebellum as a promising novel target for related research.

Another component of the RPN are the perigenual anterior cingulate cortex (pgACC), known for its integrative and modulatory role in pain processing and related decision-making^47^, and the ventral anterior insula, a region functionally connected to the pgACC and involved in emotion and disgust processing, besides its role in the processing of cognitive-affective aspects of pain^37^.

Several connections, exerting small-to-moderate influence on the prediction, involve the occipital lobe. Some of these predictive connections could be interpreted as secondary to pain perception (analogously to the well-known deactivations of visual areas during noxious stimulation) or, alternatively, might be related to the somewhat underreported effect of visual context on pain experience^48^.

### From resting-state activity to pain response

While only a few papers have previously highlighted the relationship between resting-state activity and acute pain^22,45,49^, several studies have focussed on pain anticipation^12,15–17^ i.e. on a pain-free state directly preceding a painful stimulus. In summary, results from these studies suggest that the functional state and connectivity profile of the anterior insula, periaqueductal grey, anterior cingulum, cerebellum and areas of the fronto-parietal network appear to reflect the individuals’ momentary susceptibility to potentially painful stimuli. However, from these studies it is unclear to what degree pain sensitivity is modulated by trait-like (anxiety, pain catastrophising) or by state-like (preceding emotional appraisal, attentional or pain-specific mental states) characteristics^50^. In contrast to the short periods used in anticipatory studies, our study is based on a ten-minute-long resting-state period and predicts pain sensitivity measured several days later. Thus, our results strongly support the presence of a trait-like neural signature of pain sensitivity. However, the confounder analysis (association with time between measurements and menstrual cycle) also provides evidence for temporal dynamics.

### Limitations

While the validation and testing of the proposed predictive signature is highly reliable in terms of generalizability, here we note that the applied brain atlas, while providing full-brain coverage and a generalizable functional parcellation, still introduces a-priori constrains in laterality and precise localisation of brain regions (e.g. it does not contain the PAG and other brain-stem areas).

### Outlook and implication

The identified predictive network signature has important implications from a basic research and clinical perspective and paves the way for future translational research. Investigating how the “*resting-state pain susceptibility network*” is embedded into the general resting-state brain activity could extend our knowledge about the complex functional architecture of the resting brain and foster our understanding of the mechanisms by which the subjective experience of pain emerges from neural function. Moreover, the RPN-signature allows the better characterisation of various clinical populations (e.g. in post-operative or chronic pain), especially when reliable behavioural pain reports cannot be obtained.

The RPN-signature may extend our possibilities to non-invasively characterise clinical populations (e.g. in post-operative or chronic pain), especially when reliable behavioural pain reports cannot be obtained. Our pre-registered, multicentre design covers heterogeneity originating from differences in research setting and ensures the reliability, robustness and generalizability of results necessary for this translation. A future, iterative, bootstrapping-like research process involving a stricter standardisation of experimental procedures and involving clinical populations promises to further improve the predictive capabilities of the RPN-signature.

In sum, the RPN-signature identified here has the potential to become a novel, non-invasive neuromarker for the *supraspinal neural contribution* to pain sensitivity and the susceptibility to clinical pain states, especially in translational research and development of analgesic treatment strategies, where uncoupling peripheral and central mechanisms is often of crucial interest. Moreover, the RPN-signature might serve as a novel, resting network-based building block in a future ‘pain biomarker composite signature’^24^.

## Acknowledgements

We are thankful to Katja Wiech (University of Oxford, Oxford, UK) for her valuable insights and comments on the manuscript. Funding: this research was supported by the Mercator Research Center Ruhr (MERCUR), Germany. BK was supported by the UNKP-18-3 ‘New National Excellence Program’ of the Ministry of Human Capacities, Hungary.

## Author contributions

T.S., M.Z. and U.B conceived the study, T.S. and B.K. conducted the analyses, T.S and U.B wrote the paper. *Study 1* was and performed by M.Z. and T.S-W. *Studies 2* and *3* were performed by F.S. and T.S. and by B.K. and Z.T.K., respectively. T.S., U.B., M.Z., Z.T.K., B.K. contributed to the interpretation, as well as manuscript preparation. The whole study was supervised by T.S. and U.B.

## Materials and Correspondence

Correspondence and material requests can be addressed to Tamas Spisak.

## Methods

### Study design considerations

The study design was established with careful consideration of recent recommendations, requirements and standards for neuroimaging biomarkers^1–5^ (neuromarkers) and motivated by the following thoughts.

#### Maximise predictive performance

We employed a standardized pre-processing pipeline to ensure optimal sensitivity of the neuromarker, as sufficient effect size is a basic requirement of any clinical utility^3^. We used high precision image alignment, incorporating individual anatomy when extracting fMRI time-series data. Moreover, we adopted recent recommendations^6,7^ and protocols^8^ regarding artefact reduction and optimised our workflow to meet the special needs of connectome-based analysis. We used our in-house developed, open-source python software library “Pipelines Utilising a Modular Inventory” (*PUMI*, https://github.com/spisakt/PUMI), which is based on nipype^9^, a community-based Python project providing a uniform interface to existing neuroimaging software and, in part, re-used code from the *C-PAC* ^10^ and the niworkflows^11^ open-source projects. A predictive modelling (“machine learning”) approach was utilised to exploit the rich data provided by resting-state functional brain networks and, potentially, take advantage of ‘fMRI hyperacuity’^12^.

#### Assessing predictive power under realistic conditions

Prospective validity and generalisability to new datasets is an essential property of any neuromarker candidates^3^. Internal validation techniques, often used in related research may introduce various biases when assessing predictive performance^13–15^. We therefore used a pre-registered, external validation strategy, that strictly separated model training and performance assessment. For model training, we exclusively used data from *Study 1.* We conducted two independent sub-studies (*Studies 2 and 3*) in different research centres, with different equipment and different research staff for validation. We used a “liberal” alignment of research settings, allowing for a reasonable heterogeneity in procedures, equipment, imaging sequences, language of participant-researcher communication across study-centres, introducing a reasonable heterogeneity in the validation procedure to ensure generalizability.

#### Ensure that prediction is driven by neural signal and is specific to pain sensitivity

Powerful predictions can be driven by a “third variable”, related but different from the measure in question, e.g. systematic movement artefacts. Therefore, assessing the neuroscientific validity of the predictive model is an important step in neuromarker development^3^. To ensure that the proposed marker of pain sensitivity is indeed driven by neural signals associated with pain sensitivity, we evaluated the correlation of the predicted score with various pre-defined (and pre-registered) confounder and validator variables.

#### Ensure accessibility of results

Biomarker development is a complex, multi-stage process which can be largely facilitated by transparency, accessibility and deployability of the methodology^3^. Therefore, we applied a comprehensive pre-registration and made the source code of the method open-source and freely available for the community (*https://spisakt.github.io/RPN-signature*). Moreover, we provide a platform-independent, easy to use docker container which provides the opportunity to use our predictive model as a “research product”^3^, to obtain out-of-the box pain sensitivity predictions form any appropriate imaging datasets.

### Participants

A total of N=116 healthy, young volunteers were involved in three sub-studies. Age and sex of the participants is reported in Supplementary Table S1. *Study 1* involved N_1_=39 participants (the same sample as in Ref.^16^). It was performed at the Ruhr University Bochum (Germany) by MZ and TSW and used as the training sample for the machine learning-based prediction of pain sensitivity and additionally, served as a basis for the internal validation of the prediction. *Studies 2* and *3* (N_2_=48, N_3_=29) were performed at the University Hospital Essen (Germany) by FS and TS and at the University of Szeged (Hungary) by BK and TK, respectively, and served as samples for external validation. Inclusion criteria and exclusion criteria were largely identical in all three centres and are listed in Methods Table 1. Recruitment and reimbursement policies varied across centres; participants received 20 €/h in *Studies 1* and *2* and no reimbursement in *Study 3*.

Metal implant, unremovable piercing, peacemaker, tattoo in head/neck position, pregnancy or known claustrophobia were considered as contraindication for MR measurement. Participants were required to abstain from consuming caffeine two hours before experiments (except in *Study 3*) and from consuming alcohol on the day of testing and the previous day.

The study was conducted in accordance with the Declaration of Helsinki and approved by the local or national ethics committees (Register Numbers: 4974-14, 18-8020-BO and 057617/2015/OTIG at the Ruhr University Bochum, University Hospital Essen and ETT TUKEB Hungary, respectively.) All participants gave written informed consent before testing. Imaging and quantitative sensory testing (QST) were performed on the same day in Study 1 and in average 2-3 days apart in *Studies 2* and *3 (*see Supplementary Table S1 for details). MRI measurement always preceded the QST session.

**Methods Table 1.** Inclusion and exclusion criteria during the recruitment process.

| **Inclusion criteria** | **Exclusion criteria** |
| --- | --- |
| No chronic disease | Acute or chronic neurological endocrine, or psychiatric conditions |
| Age between 18 and 40 (target: 25) | Acute infections |
| Right-handedness | Contraindication for MRI measurement |
| Non-smoking | regular medication intake (except contraceptive) |
| Equal gender distribution targeted | Recent use of psychotropic or analgesic substances |
|  | Participation in any medication-associated study in the last 6 months |
|  | Wounds, scars or any other skin conditions (e.g. neurodermitis) which could affect QST measurements on the forearm and the hands |

**Methods Table 2.** MRI scanner and sequence parameters for each centre.

|  | STUDY 1 | STUDY 2 | STUDY 3 |
| --- | --- | --- | --- |
|  | **GENERAL** |  |  |
| **SCANNER** | Philips Achieva X 3T | Siemens Magnetom Skyra 3T | **GE Discovery MR750w 3T** |
| **HEAD COIL** | 32-channel | 32-channel | 20-channel |
|  | **ANATOMICAL SCAN** |  |  |
| **WEIGHTING** | T1 | T1 | T1 |
| **SEQUENCE** | MPRAGE | MPRAGE | 3D IR-FSPGR |
| **TR** | 8500ms | 2300ms | 5.3ms |
| **TE** | 3.9ms | 2.07ms | 2.1ms |
| **RESOLUTION** | 1x1x1mm<sup>3</sup> | 1x1x1mm<sup>3</sup> | 1x1x1mm<sup>3</sup> |
| **FOV** | 256x256x220 mm<sup>3</sup> | 256x256x192 mm<sup>3</sup> | 256x256x172 |
|  | **RESTING STATE FMRI** |  |  |
| **WEIGHTING** | T2* | T2* | T2* |
| **SEQUENCE** | GE EPI | GE EPI | GE EPI |
| **TR** | 2500 ms | 2520 ms | 2500 ms |
| **TE** | 35ms | 35ms | 27ms |
| **FLIP ANGLE** | 90 | 90 | 81 |
| **PHASE ENC. DIR** | COL | A>>P | A>>P |
| **FOV** | 240x240x132 | 230x230x132 | 96x96x44 |
| **NUM. OF SLICES** | 40 | 38 | 44 |
| **SLICE THICKNESS** | 3mm | 3mm | 3mm |
| **GAP** | 0.3mm | 0.48mm | 0mm |
| **SLICE ORDER** | interleaved | interleaved | interleaved |
| **IN-PLANE RES.** | 3x3mm<sup>2</sup> | 2.45x2.45mm<sup>2</sup> | 3x3mm<sup>2</sup> |
| **ACCELERATION** | SENSE 3x | GRAPPA 2x | ASSET 2x |
| **FAT SUPPRESS** | SPiR | Fat.sat. | Fat. Sat |
| **NUM. OF VOLS** | 200 | 290 | 240 |
| **DUMMY SCANS** | 5 | 5 | 0 |
| **SCANNING TIME** | 8 min 37 sec | 12 min 11 sec | 10 min |

### Measures – functional MRI

High-resolution anatomical and open-eyed resting-state fMRI measurements were acquired from all participants. Scanning parameters (including equipment) varied across centres and are listed in Methods Table 2. During measurements, participants were instructed to lie still and relaxed, without falling asleep, and avoid any movement.

Foam padding, and in *Studies 1* and *2*, pneumatic pillows were used to restrict head movements. All anatomical MRI measurements were screened for incidental findings.

### Measures – QST

Heat (HPT), cold (CPT), and mechanical (MPT) pain thresholds were acquired according to the QST protocol^17^. Warmth (WDT), cold (CDT) and in *Study 2* and *Study 3*, mechanical (MDT) detection thresholds were obtained as additional control measures. All sensory measurements were obtained from the palmar left forearm, proximal to the wrist crest. Within the QST framework, thermal thresholds are determined using a method of limits^18^. To this end, increasing and decreasing temperatures were applied to the skin with an *MSA* thermal stimulator (Somedic, Hörby, Sweden) in Study 1 and *Pathway* thermal stimulators (Medoc Ltd., Ramat Yishai, Israel) in *Studies 2* and *3*. In all studies, ATS thermodes were used on a skin surface of 30×30mm, with a baseline temperature of 32°C. Participants were instructed to indicate the onset of pain by button press. For all thermal thresholds 6, rather than 3 (as in the original protocol)^17^, stimulus repetitions were performed to reduce between-subject variance. Furthermore, the first measurement was discarded from analysis as a test stimulus. HPT and CPT were calculated as the arithmetic means of the five remaining threshold temperatures. MPTs and MDTs were determined using a staircase method^18^. Five increasing and five decreasing trains of pinprick (MRC Systems, Heidelberg, Germany) stimuli were applied to the palmar left forearm in an alternating fashion, whereas the participant was instructed to categorize the stimuli as noxious, or non-noxious. Mechanical detection threshold was assessed analogously with von Frey filament stimulations. MPT and MDT were computed as the log-transformed geometric mean force determined in five ascending and descending staircase-thresholding-runs.

### Additional measures

Age, sex, self-reported height, weight and, for female participants, the date of the first day of the last menses and the use of contraceptives, was recorded prior to all measurements. Additionally, self-reported weekly alcohol consumption and level of education (primary school, secondary school, university) was recorder *for Studies 1 and 2*. Before the QST, participants filled out *the* Pain Sensitivity *Questionnair*e (PSQ)^19^, the Pain Catastrophizing Scale (PCS)^20^, the *State*-*Trait Anxiety Inventory* (STAI)^21,22^, the short German version of the Depression Scale (ADS-K, Center for Epidemiologic Studies)^23^ and, *additionally* in *Studies 2* and *3*, the Pittsburgh Sleep Quality Index (PSQI)^24^ and the perceived stress questionnaire (PSQ20)^25^. In *Studies 2* and *3*, blood pressure was measured both before the MRI and the QST measurements. Moreover, for Sample 1, T_50_ values were available from a parallel experiment performed on the day before fMRI testing. T_50_ represents the temperature (in °C) necessary to induce a heat-pain rating of 50 (on a scale ranging from 0, no pain to 100 unbearable pain). T_50_ values were obtained from a non-linear (2^nd^-order polynomial) interpolation of ratings obtained in response to 15 tonic heat pain stimuli (duration: 16 seconds) between 42,5°C and 48°C, presented in pseudo-randomized “grid search” fashion.

### Calculation of Pain Sensitivity

The target variable for the prediction was a single composite measure of individual pain sensitivity summarizing HPT, CPT, and MPT as defined in Ref.^16^.

In *Study 1*, HPT, CPT, and MPT were Z-transformed (mean centred and standardized) and HPT, as well as MPT were inverted (multiplied by -1), so that higher Z-values denoted higher pain sensitivity. Then, the arithmetic mean of the Z-transformed variables was computed for each participant and defined as “pain-sensitivity score”. In *Studies 2* and *3*, the same procedure was applied, except that Z-transformation was based on the population-mean and standard deviation of *Study 1*, to ensure that the same scale was used across studies. Extreme QST values were defined using the normative 95% percentiles reported in Ref.^17^; participants showing extreme HPT, CPT or MPT values in at least two of the three modalities were excluded. This screening resulted in excluding 0, 3 and 2 participants in *Samples 1, 2* and *3*, respectively (Table S2).

### fMRI preprocessing

As fMRI-based functional connectivity is susceptible to in-scanner motion artefacts^6–8,26,27^, appropriate preprocessing and signal cleaning is key to successful connectivity-based prediction. Resting-state functional MRI data was preprocessed identically in all three studies. The applied, nipype-based workflow is depicted on Figure S1. It utilised third-party neuroimaging software, code adapted from the software tools *C-PAC*^10^ and *niworkflows*^11^, and in-house *python* routines.

Brain extraction from both the anatomical and the structural images, as well, as tissue-segmentation from the anatomical images was performed with *FSL bet* and *fast*^28^. Anatomical images were linearly and non-linearly co-registered to the 1mm-resolution MNI152 standard brain template brain with *ANTs*^29^ (see https://gist.github.com/spisakt/0caa7ec4bc18d3ed736d3a4e49da7415 for source code). Functional images were co-registered to the anatomical images with the boundary-based registration technique of *FSL flirt*^28^. All resulting transformations were saved for further use. The preprocessing of functional images happened in the native image space, without resampling. Realignment-based motion correction was performed with *FSL mcflirt*^28^. The resulting six head motion estimates (3 rotations, 3 translations), their squared versions, their derivates and the squared derivates (known as the Friston-24-expansion^30^) was calculated and saved for nuisance correction. Additionally, head motion was summarised as frame-wise displacement (FD) timeseries, according to Power’s method^26^, to be used in data censoring and exclusion. After motion-correction, outliers (e.g. motion spikes) in time series data were attenuated using *AFNI despike*^31^. The union of the eroded white-matter maps and ventricle masks were transformed to the native functional space and used for extracting noise-signal for anatomical CompCor correction^32^.

In a nuisance regression step, 6 *CompCor* parameters (the 6 first principal components of the noise-region timeseries), the Friston-24 motion parameters and the linear trend were removed from the timeseries data with a general linear model. On the residual data, temporal bandpass filtering was performed with *AFNI’s 3DBandpass* to retain the 0.008-0.08Hz frequency band. The prior use of *AFNI’s despike* is expected to attenuate aliasing of residual motion artefacts into the neighbouring time-frames during bandpass filtering^33–35^. To further attenuate the impact of motion artefacts, potentially motion-contaminated time-frames, defined by a conservative FD>0.15mm threshold, were dropped from the data (known as “scrubbing” the data)^33^. Participants were excluded from further analysis if the mean FD exceeded 0.15mm, or when more then 30% of frames were “scrubbed”. This resulted in exclusion of 4, 8 and 7 participants in *Samples 1, 2* and *3*, respectively (Table S2). Quality-control (registration-check, carpet-plots, see e.g. Figures S2-4) was performed throughout the workflow.

### Functional connectivity analysis

The 122-parcel version of the *MIST* ^36^ multi-resolution functional brain atlas and grey matter masks obtained from the anatomical image were transformed to the native functional space. Native-space atlas regions were then masked with the grey matter masks that were obtained from the anatomical image and transformed to functional space previously. Voxel-timeseries were averaged over these individualised MIST regions and, together with the mean grey matter signal, retained for graph-based connectivity analysis.

Regional timeseries were ordered into large-scale functional modules (defined by the 7-parcel *MIST* atlas) for visualization purposes (Figure 1). Partial correlation was computed across all pairs of regions (and global grey matter), as implemented in the *nilearn*^37,38^ python module. Partial, rather than simple correlations were used to rule out indirect connectivity^39^. Our graph-modelling approach ensured that the global-grey matter signal is handled as a confound during computing the partial correlation coefficients but, at the same time, also considered it as a signal of interest, as it may represent vigilance related processes^40^. Partial correlation coefficients were organised to 123 by 123 (122 regions + global grey matter signal) symmetric connectivity matrices. The upper triangle of these matrices was used as the feature space for machine learning-based predictive modelling.

### Predictive model training and validation

Whole-brain resting-state functional connectivity data of *study 1* (*N*_*1*_*=35*, after all exclusions ^16^, Supplementary Table S2) was used as the input feature-space (*P=7503* features per participant) to predict individual pain sensitivity scores, leading to a “large *P* — small *N*” setting.

We constructed a machine learning pipeline (https://github.com/spisakt/RPN-signature/blob/master/PAINTeR/model.py) in *scikit-learn*^37,41^, consisting of robust feature scaling (removes the median and scales with data quantiles), pre-selection of features^42^, selecting the *K* “best” features with strongest relationships to the target variable and an Elastic Net regression model43 (a linear model with combined *L*_*1*_ and *L*_*2*_-norms as regulariser). Free hyperparameters of the machine learning pipeline were the number of pre-selected features (*K*), the ratio of the *L*_*1*_*/L*_*2*_-regularization and the weight (alpha) of regularisation. Hyperparameters were optimised with a grid-search procedure and negative mean squared error as cost function. Values for *K* ranged from 10 to 200 with increments of *5*, and included [.1, .5, .7, .9, .95, .99, .999] for the *L*_*1*_*/L*_*2*_ ratio [.001, .005, .01, .05, .1, .5] for alpha. Hyperparameter optimisation was performed in a leave-one-participant-out cross validation (internal validation phase). Cross-validation incorporated the complete machine-learning pipeline to avoid introducing dependencies between the training and test samples. Note that fMRI preprocessing was independent between subjects, thus not included in the cross-validation. Optimal hyperparameters were found to be *K=25, L*_*1*_*/L*_*2*_*-ratio=0.999* and alpha=0.005.

External validation was performed by applying the RPN-signature on the fMRI data of *Studies 2* and *3 (N*_*2*_*=37, N*_*3*_*=19*, after exclusions, Supplementary Table S2). The resulting predictions were compared with the observed QST-based pain sensitivity scores by calculating mean squared error and explained variance. In addition, we fitted a linear model to predicted and observed pain sensitivity scores, obtaining p-values for the relationship.

### Confounder analysis

To explore potential confounding variables, the predicted pain sensitivity-scores (or cross-validated predictions in case of *Sample 1*) were contrasted to mean and median FD, the percentage of scrubbed volumes, systolic and diastolic blood pressure before both the MRI and QST measurement (as blood pressure was earlier reported^44^ to be associated with sensitivity to mechanical pain), the time delay between MRI and QST testing (to test for temporal stability of the prediction), age, sex, BMI, number of days since the first day of the last menses, alcohol consumption (units/week), level of education, state and trait anxiety (STAI), score of depressive symptoms (ADS-K), self-reported pain sensitivity (PSQ) and pain catastrophising (PCS), perceived stress (PSQ20), quality of sleep (PSQI), and non-noxious QST detection thresholds (CDT, WDT and MDT, where available). Moreover, in Study 1 predictions were compared to T50-values and MR spectroscopy-based GABA and Glutamate/Glutamine levels in pain-processing brain regions (see Ref.^16^ for details). Associations were tested with linear models.

### Visualization of the predictive network

The predictive interregional connections highlighted by the non-zero regression coefficients of the RPN-signature were displayed as a ribbon plot using the R-package *circlize*^45^ (Figure 3). Corresponding individualised brain region masks were transformed back to standard space to create a study-specific regional probability map (reflecting co-registration accuracy and individual variability in morphology). Probability maps were multiplied by the sum of corresponding regression coefficients to create a “regional predictive strength” map, which was then visualised with *FSLeyes*^28^ and *MRIcroGL* (https://www.mccauslandcenter.sc.edu/mricrogl) (Figure 3). Overlap of the regional predictive strength map with two other pain-related maps was also visualized with *FSLeyes* and *MRIcroGL* (Figure 4). The analysis of large-scale resting-state network-involvement (as defined by the *MIST* ^36^ brain atlas) was performed by summarising and Z-transforming the voxel values across the seven regions-of-interest. Polar plot was made with the R-package *ggplot2*^46^.

### Data and software availability

Processed data (regional timeseries) and source code are deposited at https://github.com/spisakt/RPN-signature. Raw imaging data is available at openneuro.org. The RPN-signature scores can be computed based on structural and resting-state functional datasets by the software tool with the same name. The RPN-signature software tool consists of the described MRI processing pipeline and the functional connectome-based predictive model. It is available as source code at https://github.com/spisakt/RPN-signature. As the software follows the Brain Imaging Data Structure (BIDS)^47^ and the BIDS-App specification, it provides a standard command line interface and relies on Docker-technology. The docker image is deposited on Docker Hub: (https://cloud.docker.com/repository/docker/tspisak/rpn-signature) and does not depend on any software outside the container image. This, together with the fully transparent continuous integration-based development and automated tagging and versioning, enhances software availability and supports reproducibility of RPN-signature results.

## Supplementary Materials

**Table S1.** Summary statistics of pain sensitivity, in-scanner motion and demographic data. Summary statistics were computed after exclusions. Abbreviations: BMI: body mass index, FD: framewise displacement, Diff MRI-QST: difference between MRI and QST measurements in days, PSQ: pain sensitivity questionnaire, PCS: pain catastrophising questionnaire, STAI: state-trait anxiety inventory, PSQI: Pittsburgh sleep quality index, ADS-K: German depression scale, PSQ20: perceived stress questionnaire, QST: quantitative sensory testing, pain sens.: pain sensitivity, MAE: mean absolute error.

|  | *Study 1* | *Study 2* | *Study 3* |
| --- | --- | --- | --- |
| **Demographics** |  |  |  |
| Site location | *Bochum, GER* | *Essen, GER* | *Szeged, HUN* |
| N before exclusion | 39 | 48 | 29 |
| N after exclusion | 35 | 37 | 19 |
| Age (mean ± sd) | 26.1 ± 3.9 | 24.9 ± 3.5 | 24.8 ± 3.1 |
| Age (min, max) | 20, 39 | 19, 36 | 21, 31 |
| Sex (%female) | 37% | 54% | 53% |
| BMI (mean ± sd) | 23.4 ± 2.3 | 23.2 ± 2.7 | 23.1 ± 4.3 |
| **Confounders - MRI** |  |  |  |
| meanFD (mean ± sd) | 0.08 ± 0.02mm | 0.07 ± 0.02mm | 0.07 ± 0.02mm |
| meanFD (min, max) | 0.04, 0.13 | 0.04, 0.12 | 0.04, 0.12 |
| %scrubbed (mean ± sd) | 7.1 ± 7.4% | 6.4 ± 6.7% | 8.4 ± 8.0% |
| %scrubbed (min, max) | 0, 26% | 0, 29% | 2, 26% |
| Diff MRI-QST (mean ± sd) | 0 ± 0 days | 3.1 ± 2.9 days | 2.3 ± 1.4 days |
| Diff MRI-QST (min, max) | 0, 0 days | 1, 18 days | 1, 5 days |
| **Confounders - questionnaires** |  |  |  |
| PSQ pain sensitivity (mean ± sd) | 56.6 ± 19.1 | 55.3 ± 18.5 | 44.7 ± 14.8 |
| PCS catastrophising (mean ± sd) | 15.47 ± 7.9 | 14.9 ± 9.6 | 9.8 ± 10.4 |
| STAI state anxiety (mean ± sd) | 42.8 ± 4.0 | 33.4 ± 7.4 | 31.6 ± 10.2 |
| STAI trait anxiety (mean ± sd) | 44.4 ± 3.1 | 37.2 ± 9.9 | 31.2 ± 8.6 |
| PSQI sleep quality (mean ± sd) | N/A | 6.1 ± 3.1 | 3.0 ± 2.2 |
| ADS-K depression (mean ± sd) | 7.0 ± 4.1 | 7.8 ± 6.6 | 5.8 ± 5.9 |
| PSQ20 stress (mean ± sd) | N/A | 44.4 ± 9.4 | 26.5 ± 4.0 |
| **Prediction target** |  |  |  |
| QST pain sens. (mean ± sd) | 0.008 ± 0.73 | 0.29 ± 0.80 | -0.44 ± 0.46 |
| QST pain sens. (min, max) | -1.45, 1.53 | -1.82, 1.57 | -1.2, 0.56 |
| **Prediction** |  |  |  |
| MAE | 0.271 | 0.553 | 0.248 |
| MSE | 0.316 | 0.536 | 0.173 |
| Explained variance | 39.2% | 18.1% | 17.0% |
| Pearson's r | 0.63 | 0.43 | 0.47 |
| p-value | 0.00006 | 0.008 | 0.04 |

**Table S2.**
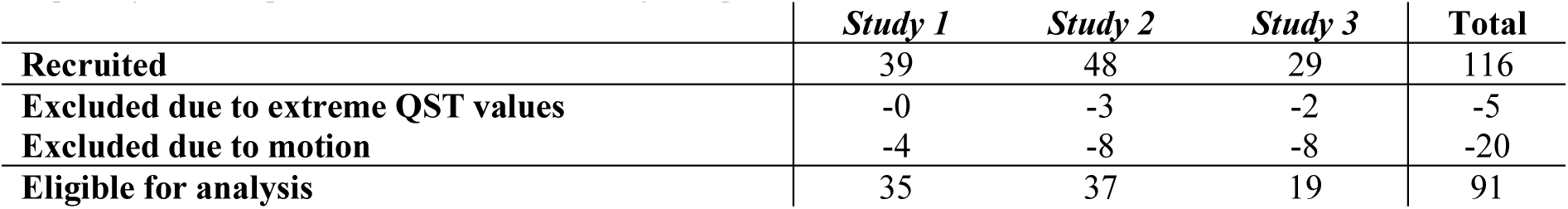
Recruitment numbers and exclusions per criteria. *For privacy reasons, possible cases with incidental findings are not shown in the table.*

|  | ***Study 1*** | ***Study 2*** | ***Study 3*** | **Total** |
| --- | --- | --- | --- | --- |
| **Recruited** | 39 | 48 | 29 | 116 |
| **Excluded due to extreme QST values** | -0 | -3 | -2 | -5 |
| **Excluded due to motion** | -4 | -8 | -8 | -20 |
| **Eligible for analysis** | 35 | 37 | 19 | 91 |

**Table S3.** Results of confounder analysis not shown in main Table 1. Trait anxiety and blood pressure (systolic and diastolic) as measured on the day of QST measurement.

|  | **anxiety trait** |  | **BP sys. QST** |  | **BP dias. QST** |  |
| --- | --- | --- | --- | --- | --- | --- |
|  | **R<sup>2</sup>** | **p** | **R<sup>2</sup>** | **p** | **R<sup>2</sup>** | **p** |
| ***Study 1*** | 0.000 | 1.000 | N/A | N/A | N/A | N/A |
| ***Study 2*** | 0.001 | 0.876 | 0.010 | 0.547 | 0.002 | 0.816 |
| ***Study 3*** | 0.040 | 0.412 | 0.013 | 0.723 | 0.110 | 0.291 |

**Figure S1.**
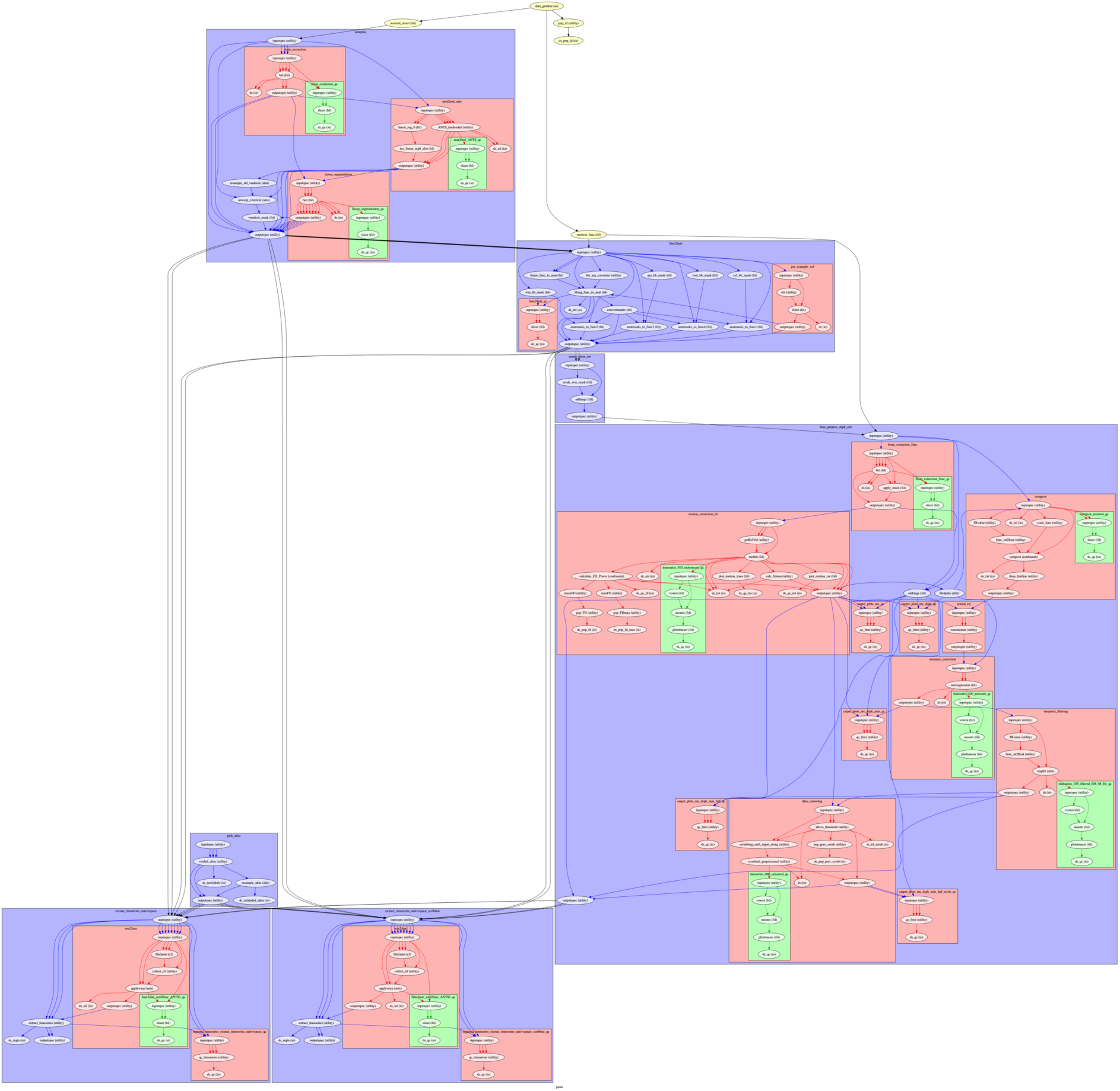
Nipype workflow of the MRI image preprocessing and connectivity calculation for the RPN-signature. Workflow is based on the in-house PUMI workflow library system (*https://github.com/spisakt/PUMI*). Source code of the workflow: *https://github.com/spisakt/PAINTeR/blob/master/pipeline/pipeline_PAINTeR.py* High resolution workflow graph is available on-line: *https://github.com/spisakt/PAINTeR/blob/master/pipeline/graph.png*

**Figure S2.**
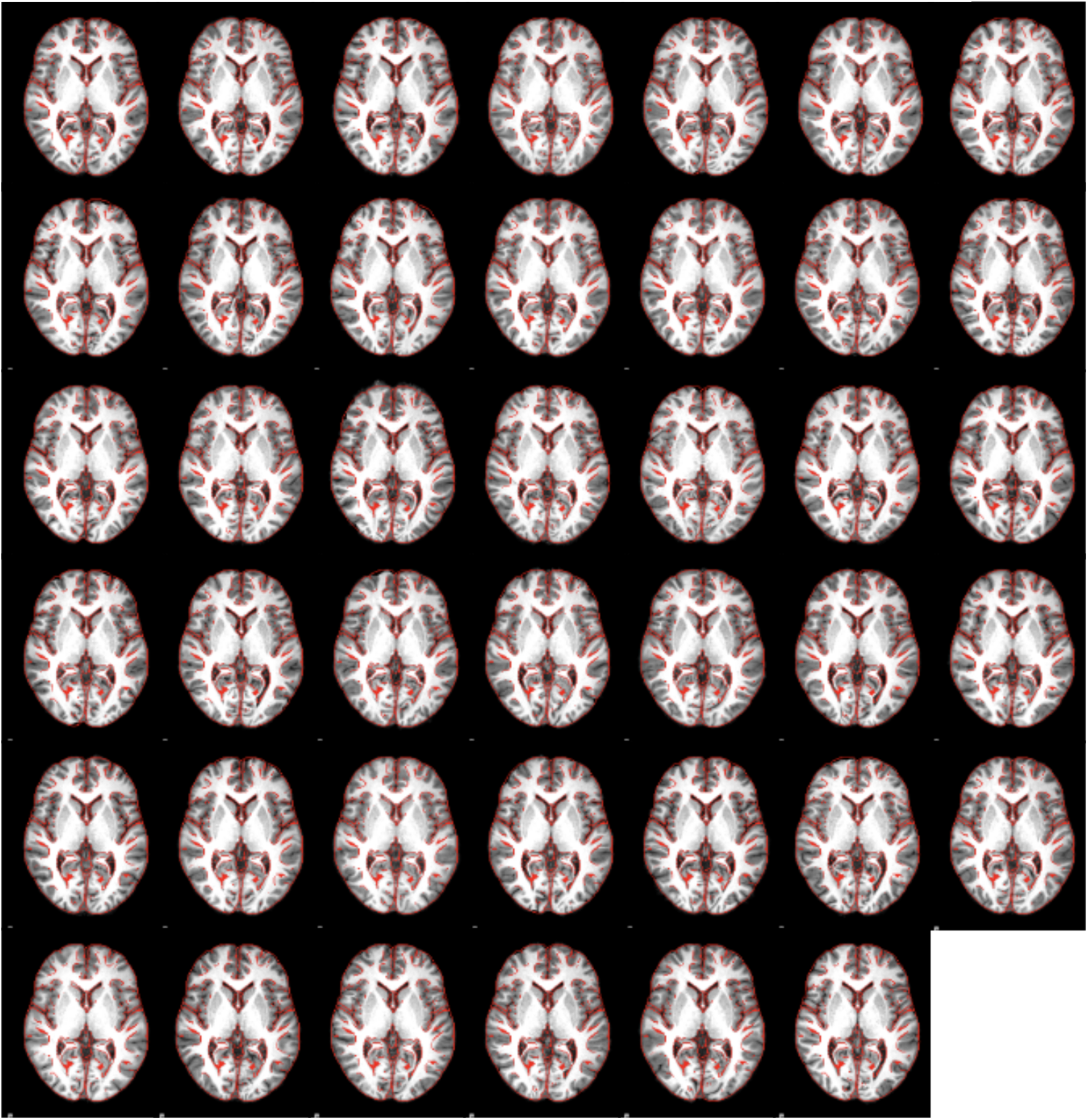
Quality-check images for the spatial standardisation of the anatomical images in Study 1.

**Figure S3.**
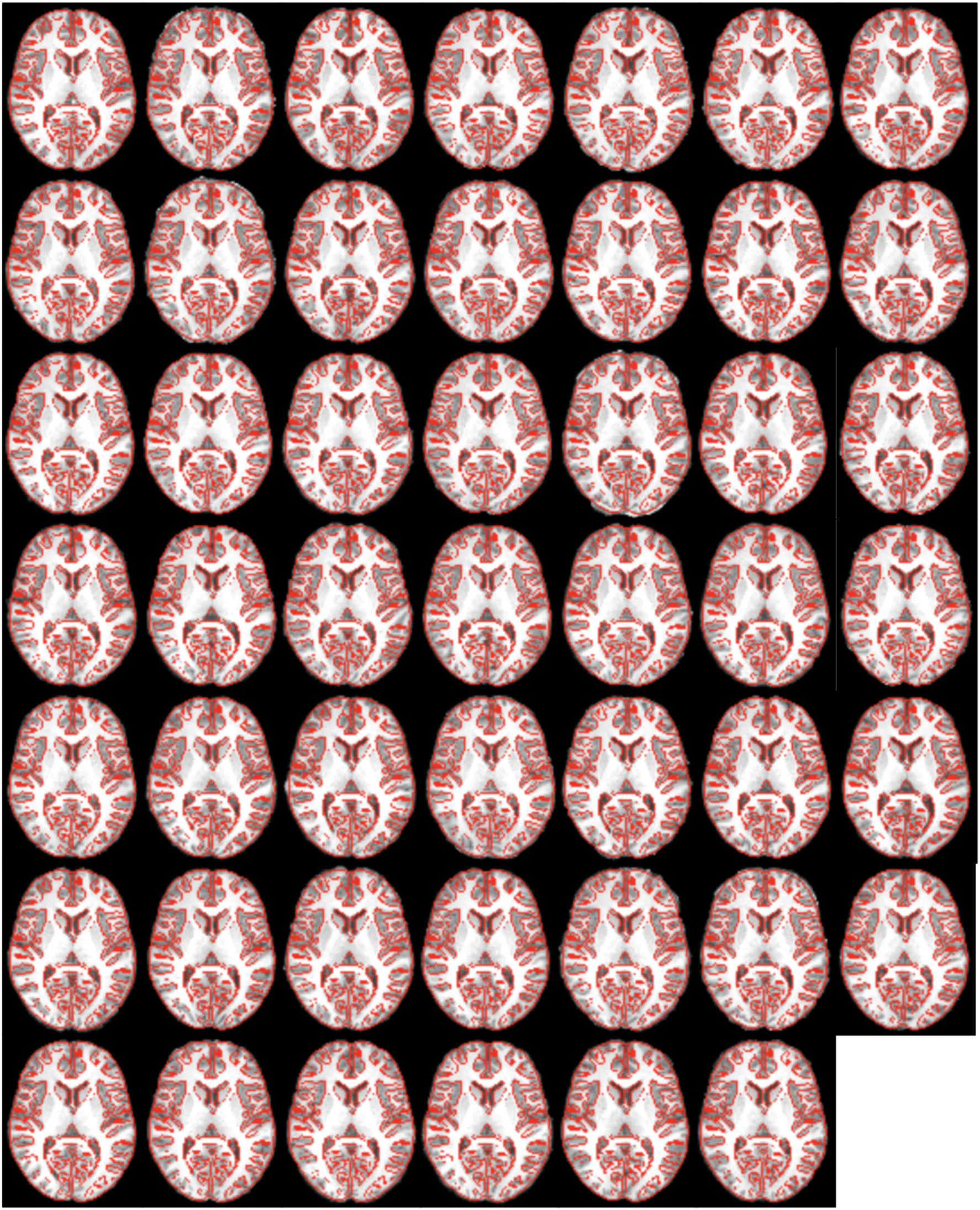
Quality-check images for the spatial standardisation of the anatomical images in Study 2.

**Figure S4.**
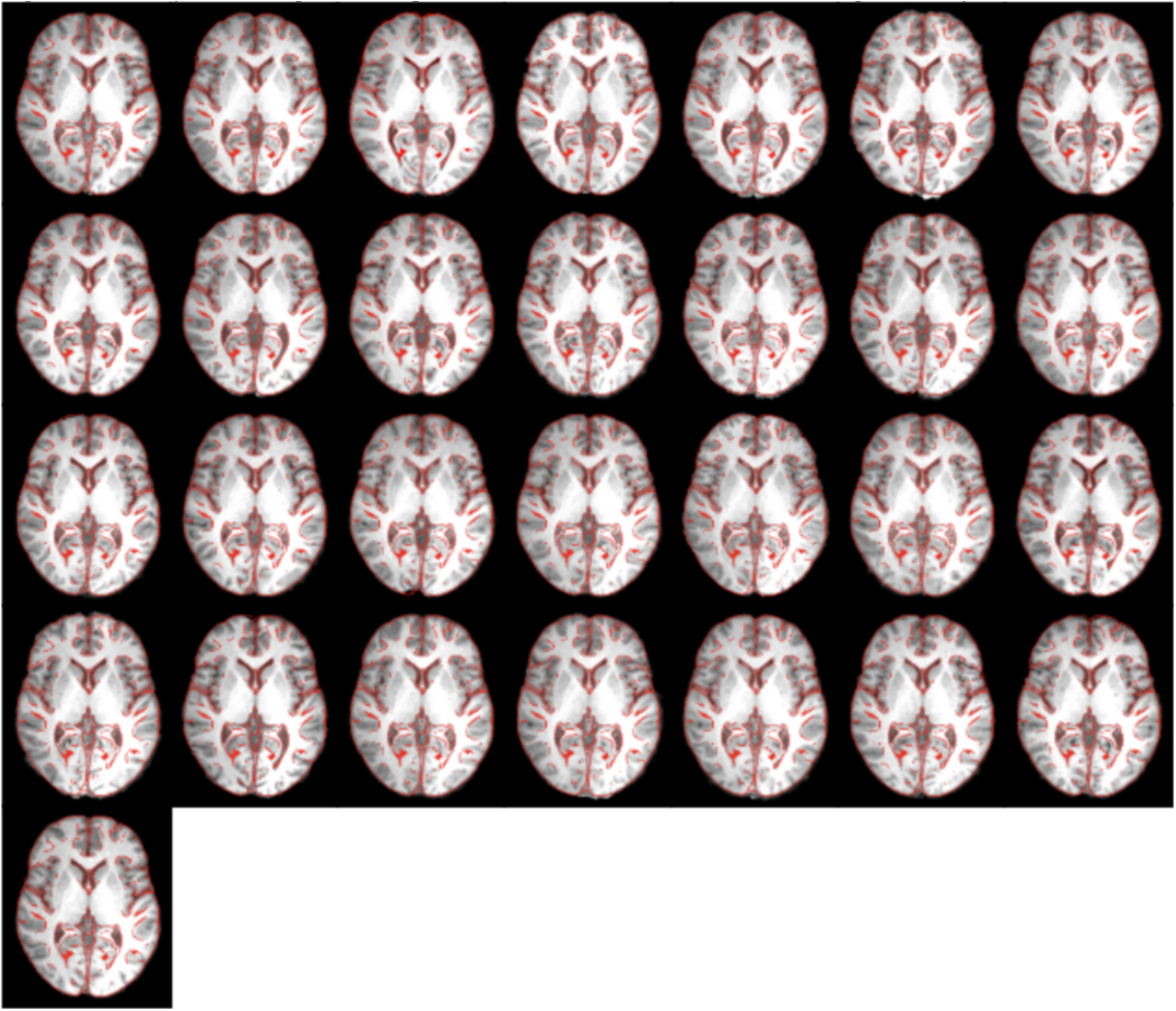
Quality-check images for the spatial standardisation of the anatomical images in Study 3.

